# Spinal electrophysiology reveals frequency-specific spatial patterns of neural activity and corticospinal coherence during pincer-grip

**DOI:** 10.1101/2025.11.11.687864

**Authors:** Prabhav Mehra, Saroj Bista, Marjorie Metzger, Eileen R. Giglia, Serena Plaitano, Leah Nash, Éanna Mac Domhnaill, Peter Bede, Muthuraman Muthuraman, Orla Hardiman, Madeleine Lowery, Bahman Nasseroleslami

## Abstract

**Background:** Neural dynamics within sensory-motor networks involved in motor control exhibit well-established frequency-dependent patterns of cortical activity. In contrast, corresponding neural patterns of spinal cord activity remain poorly understood. As an active functional part of motor control, characterising spinal cord activity is essential for understanding sensorimotor function. High-density electrospinography presents a novel technique to assess spinal neural dynamics by non-invasively recording task-relevant electrical activity.

**Objective:** To non-invasively investigate spatio-spectral patterns of task-related spinal activity and its functional connectivity with cortical regions during isometric pincer-grip contraction.

**Methodology:** Here, we simultaneously recorded 128-channel electroencephalography (EEG), 64-channel electrospinography (ESG), and two bipolar electromyography (EMG) signals during an isometric pincer-grip task. Frequency-specific spatial patterns of spinal activity and cortico-spinal connectivity were evaluated by calculating task-related ESG power and cortico-spinal coherence between ESG and EEG signals during the isometric hold.

**Results:** Distinct frequency-specific spatial patterns of spinal activity were observed at lower cervical levels during sustained hold, with significant activation over the ipsilateral anterolateral region. In the beta band, task-related spinal activity was significantly ipsilateralised, and exhibited significant cortico-spinal connectivity between contralateral motor cortex and ipsi-anterolateral spinal region.

**Conclusions:** This first-of-its-kind application of HD non-invasive spinal electrophysiology revealed that the spinal cord exhibits distinct frequency-specific spatial activation and cortical connectivity patterns during pincer-grip sustained hold. Specifically, the ipsi-anterolateral spinal region demonstrated high task-relevance, potentially indicating anterior horn activity and its connectivity with motor cortex. Furthermore, beta-band activation was observed as a key signature during sustained hold, further underpinning its relevance during motor control.

## Introduction

Understanding the neural mechanisms underlying motor control is fundamental for evaluating function and dysfunction in health and disease. While non-invasive electrophysiological and neuroimaging methods played a major role in uncovering the cortical basis for motor control, the contribution of the spinal cord has largely been neglected and is often treated as a passive relay between the brain and periphery. However, multiple findings observe that the spinal cord plays an active role during motor control, including processing, transmitting, and functional integration of sensorimotor signals (Hochman, 2007; Nielsen, 2004; Pierrot-Deseilligny and Burke, 2012; Wolpert et al., 2011). Stachowski and Dougherty further highlight the importance of spinal circuits in modulating both descending and ascending information by integrating sensorimotor signals at the spinal level (Stachowski and Dougherty, 2021). Additionally, recent studies suggest the potential role of the spinal cord in motor learning (Bruel et al., 2024; Vahdat et al., 2015). Together, these findings emphasise the need to consider spinal activity as an integral component of the motor system.

Traditionally, techniques such as transcranial magnetic stimulation (TMS) and peripheral stimulation have been utilised to indirectly assess spinal cord function (de Carvalho and Swash, 2016; Fatehi et al., 2018; Fernández, 2021; Šoda et al., 2023), by probing either descending or ascending pathways. The reliance of these stimulation-based electrophysiology methodologies on evoked or reflex responses limits our ability to investigate neural complexities of the spinal cord, failing to capture the essential sensory-motor interactions during voluntary control. At the cortical level, direct electrophysiology techniques have been utilised to reveal frequency-dependent neurophysiological function during motor control (Baker, 2007; Muthukumaraswamy, 2010; Pfurtscheller et al., 2003). In particular, beta band activity has been observed as a characteristic neural signature of sustained contraction and is broadly associated with promoting existing contraction and descending control (Baker, 2007; Chakarov et al., 2009; Engel and Fries, 2010; Kilner et al., 1999). Similarly, beta-band cortico-muscular coherence (CMC) has been widely observed as a functional signature of sustained contraction (Bao et al., 2021; Bista et al., 2023; Coffey et al., 2021; Conway et al., 1995), using electroencephalography (EEG) and electromyography (EMG) signals. These studies suggest distinct spatio-spectral neuro-dynamics during motor control (Crone et al., 1998a, 1998b). However, such studies largely overlook spinal contribution, leaving out a key component within the sensorimotor pathway.

The lack of direct spinal neurophysiology studies can be attributed to significant challenges presented by the spinal cord’s complex structure and location in recording functional neurophysiological signals (Bede et al., 2012; Kinany et al., 2023). These challenges are further amplified during motor control tasks, due to increased physiological noise (muscle) and movement artifacts. Recent spinal functional magnetic resonance imaging (fMRI) studies, leveraging technological advancements (De Leener et al., 2018, 2017; Yacoub and Wald, 2018), have revealed localised spinal activation over C5-T1 vertebral regions during upper-limb voluntary motor tasks (Kinany et al., 2019; Landelle et al., 2021). However, findings regarding the lateralisation of spinal activity are inconsistent, with reports of both ipsilateral (Weber et al., 2016) and bilateral activation patterns (Bouwman et al., 2008). Moreover, because MRI is an indirect measure (neural correlate) of neural activity with low temporal resolution, it limits its capacity to study frequency-dependent neural dynamics observed during sensorimotor communication. High-density spinal neurophysiology techniques, such as electrospinography (ESG) (Bailey et al., 2024; Chander et al., 2022; Nierula et al., 2024) and magnetospinography (Akaza et al., 2020; Sumiya et al., 2017), have the potential to bridge this gap, but these investigations have largely been limited to studying evoked spinal responses. Recently, a feasibility study by Steele et al. demonstrated that non-invasive ESG can be utilised to study spinal neural dynamics during voluntary lower-limb movement (Steele et al., 2024). We recently developed a standardised electrode system to facilitate reproducible and spatially-defined recording of high-density ESG (Mehra et al., 2025), which can be utilised to investigate signatures of spinal neural dynamics at the upper spinal cord during motor tasks.

Here, we aim to characterise the spectral and spatial features of spinal electrical activity and its interaction with the motor cortex during voluntary upper limb motor control. Utilising a novel multimodal high-density ESG and EEG recording approach combined with advanced analytical methods, task-relevant spinal activation patterns and cortico-spinal coherence were investigated during sustained pincer-grip contraction. This work seeks to characterise neurophysiological signatures of spinal activity during voluntary motor control, advancing our understanding of multisystem (cortico-spinal) neural dynamics.

## Results

We recorded 64-channel high-density electrospinography (ESG) signals alongside 128-channel high-density electroencephalography (EEG) and bipolar electromyography signals from first dorsal interosseous (FDI) and abductor pollicis brevis (APB), to investigate the neural activity patterns of the upper spinal cord and its connectivity with the motor cortex during isometric motor control. During the experiment, each participant (n=10, age: 28 ± 4, 4M) performed 30 sessions of the isometric pincer-grip task at 10% of their maximum voluntary contraction (MVC). Each session lasted 15s, consisting of three guided phases: rest phase (5s), followed by isometric hold phase (5s), followed by relaxation phase (5s) (Figure 1).

**Figure 1:**
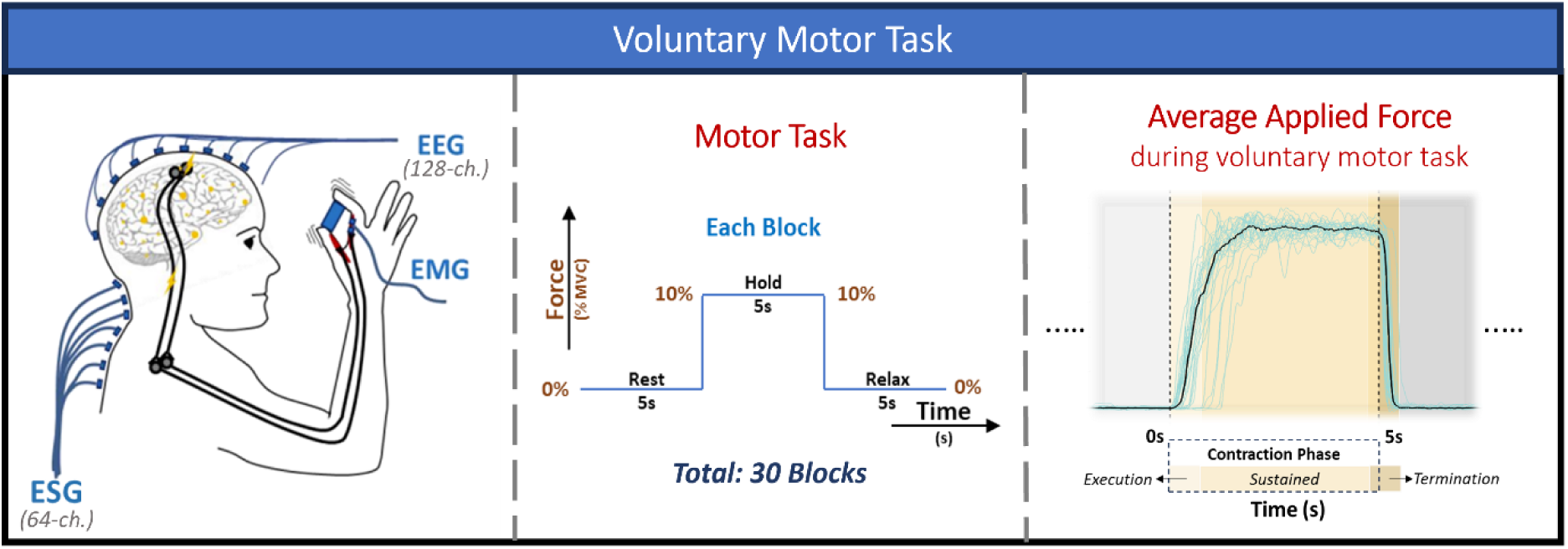
Schematic representation of the recorded signals and the voluntary motor task. Voluntary motor task consisted of three guided phases: rest phase (5s), isometric hold phase (5s), relaxation phase (5s).

### Spatial Activation Patterns of the Spinal Cord During Sustained Contraction: Frequency-band specific spectral analysis

To investigate the spatial patterns of spinal cord neural activity during pincer-grip hold, we identified task-relevant ESG channels that showed significant activation during sustained contraction compared to baseline. ESG channels exhibiting statistically significant differences in average spectral power during sustained contraction compared to baseline spectral power were classified as task-relevant. Topographical map of the baseline-corrected spectral power for the identified task-relevant channels, referred to as task-relevant power (pow_task_), revealed significant spinal activity over C5-C7 cervical vertebral levels. This activation over lower cervical vertebral levels (C5-C7) was expected, considering it corresponds to the anatomical location of motoneurons innervating the finger and thumb muscles.

### Frequency-band Specific Spatial Patterns

Next, we investigated if spinal neural activity demonstrated distinct spatio-spectral characteristics, i.e. frequency-dependent spatial activation patterns. For this, task-relevant channels were identified in three frequency bands: beta (13-30Hz), low-gamma (30-60Hz), and high-gamma (60-90Hz). Band-specific pow_task_ for the resulting task-relevant channels for each frequency band was plotted over the SC10X/U electrodes (Figure 2), to identify underlying frequency-band-specific spatial activation patterns of spinal cord activity during sustained hold.

**Figure 2:**
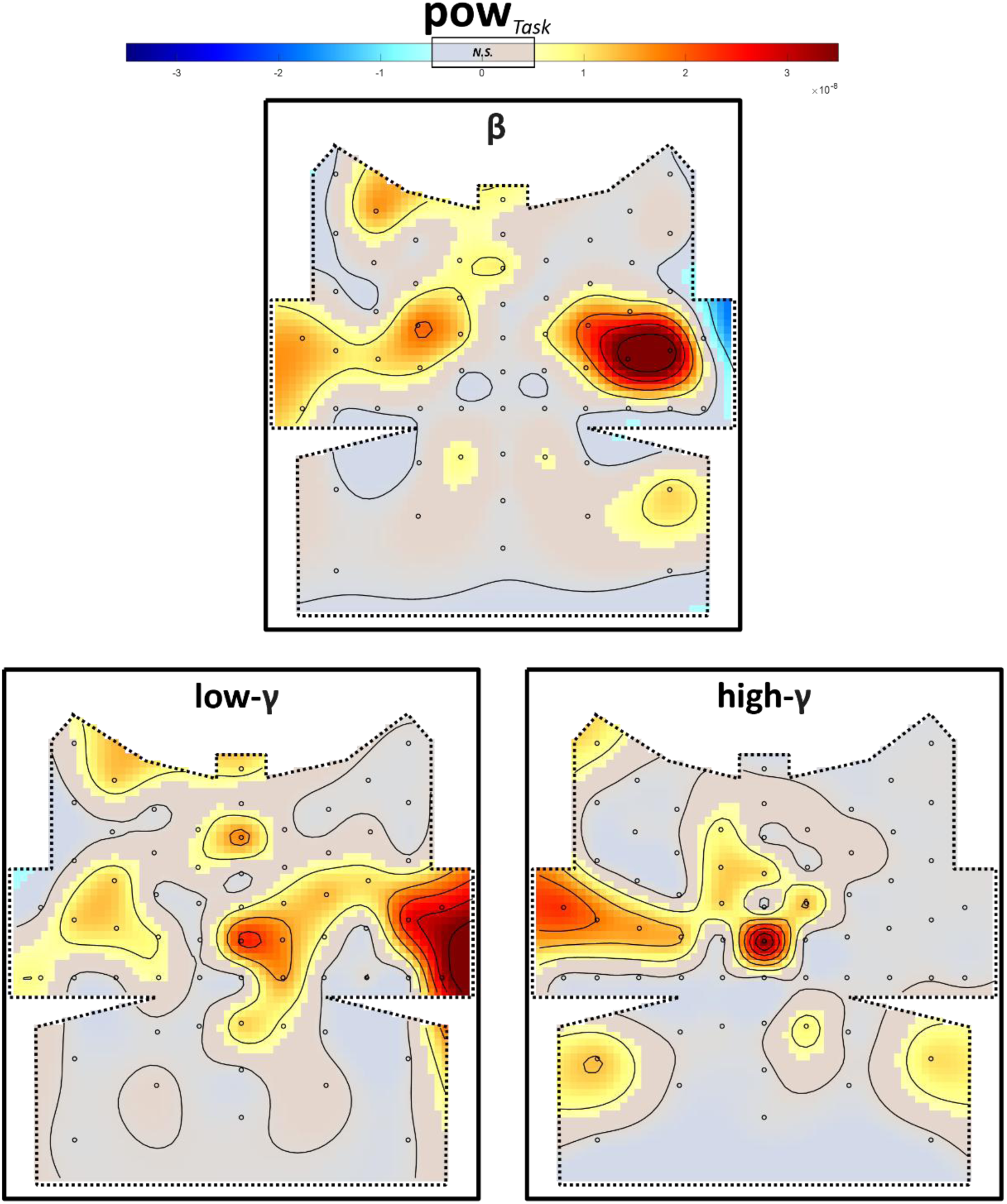
Topography maps of task-relevant power are shown for beta, low-gamma, and high-gamma frequency bands. Band-specific pow_task_ represents spectral power (baseline-corrected) during sustained contraction, for statistically different spectral power as compared to baseline (during rest). Beta-band and low-gamma band power showcased significant ipsilateral activation. In addition to ipsilateral activation, significant task-related activation at dorsal-midline (ML6) was observed in the low-gamma band. No significant ipsilateral activity was observed for high-gamma-band pow_task_ but significant dorsal-midline activation at ML6 was observed. [N.S. refers to not significant]

Distinct frequency-band-specific spatial activation patterns were identified at C5-C7 vertebral levels, as shown in Figure 2. Task-relevant beta-band ESG activity showed a dominant ipsilateral activation pattern, with its epicentre at the LL8 and LP8 electrodes and no dorsal activation. In contrast, low-gamma-banded task-relevant ESG activation was observed over both ipsilateral and dorsal-midline channels. Meanwhile, only dorsal-midline ESG activation at ML6 electrode (C6 vertebral level) was observed for high-gamma-band specific activity.

### Lateralisation

We investigated whether spinal activity demonstrated band-specific lateralisation during sustained contraction. To assess band-specific lateralisation, we compared the percentage change in ESG band power (task relative to rest) between the ipsi-anterolateral and contra-anterolateral regions. The ipsi-anterolateral region comprised LL8 and AL4 channels, while the contra-anterolateral region comprised the corresponding LL8 and AL4 channels on the opposite side, as shown in Figure 3.

**Figure 3:**
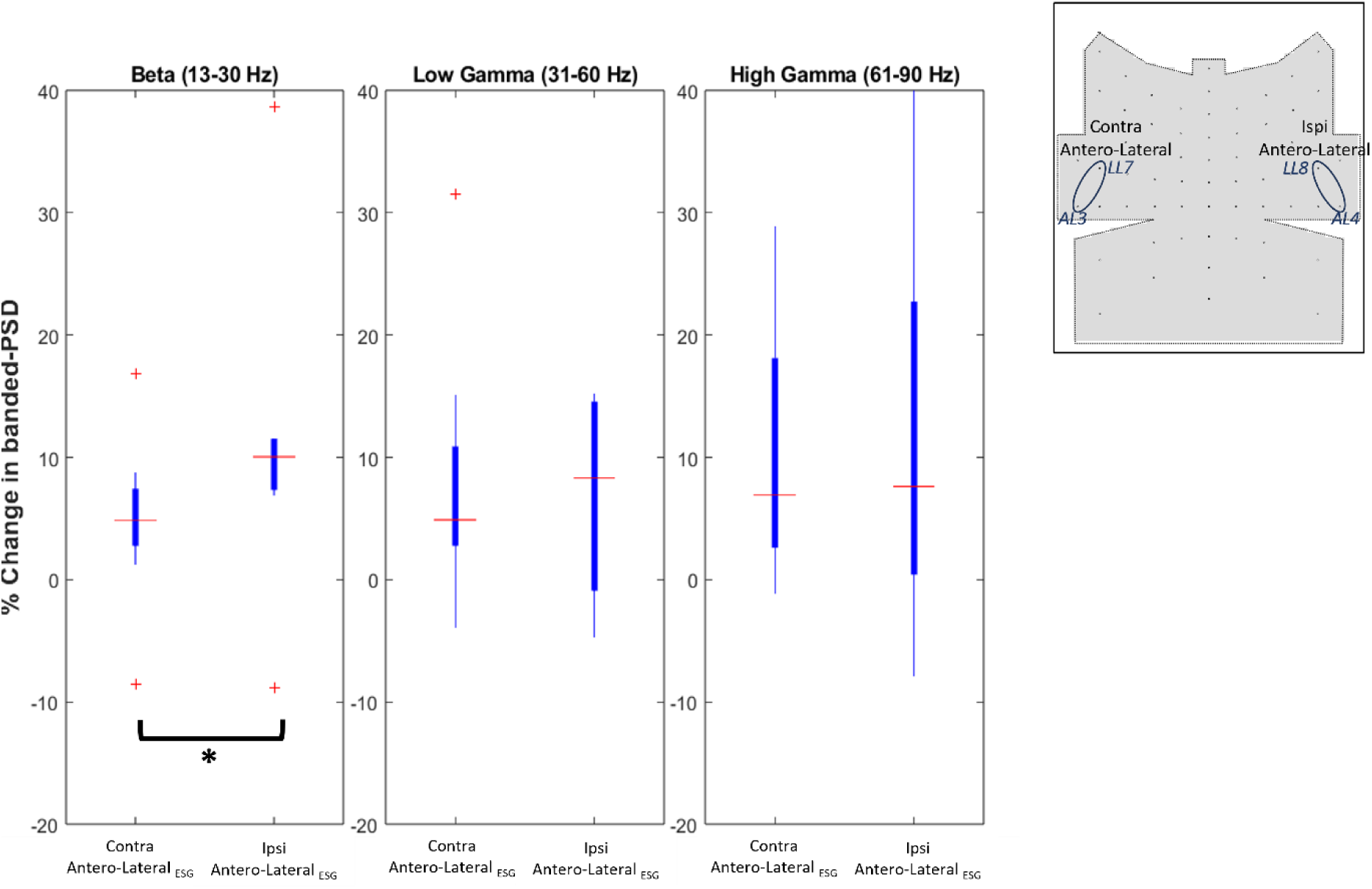
Distribution plot (box plot) of the spectral power as percentage change (task spectral power compared to baseline spectral power) for each frequency band and two ESG regions: Contra-anterolateral and Ipsi-anterolateral, as indicated on the topography map. For the beta band, a significant difference between contra-anterolateral and Ipsi-anterolateral spectral power was observed. No significant difference between the two regions was observed for low-gamma and high-gamma band, indicating beta band-specific lateralized activation during sustained hold.

A beta-band specific lateralisation of spinal activity was observed during sustained hold (Figure 3), i.e., a significant increase (p = 0.0156) in spectral power was observed at ipsi-anterolateral electrodes compared to the contra-anterolateral electrodes. In contrast, no significant differences were observed between the two regions in the low-gamma and high-gamma bands, indicating no lateralisation of spinal activity in these bands. These findings reveal frequency-specific lateralisation characteristics of spinal activity at lower cervical levels, with only beta-band activity indicating significant lateralisation.

### Functional connectivity between primary motor cortex and lower cervical spinal cord: Cortico-spinal coherence analysis

In order to examine cortico-spinal functional connectivity, coherence was estimated between EEG (C3: contralateral primary motor region, C4: ipsilateral primary motor region) and ESG at lower cervical regions (ipsi-anterolateral (LL8, AL4), contra-anterolateral (LL7, AL3)), as shown in Figure 4a. EEG recorded over the contralateral primary motor region (C3) exhibited significant coherence with ESG signals recorded over the ipsi-anterolateral spinal region in the beta band (Figure 4a). No significant coherence was detected between any other combinations of EEG and ESG signals in the beta band, revealing location-specific beta-band cortico-spinal connectivity. The location-specific beta-band coherence can be attributed to long-range connectivity via the corticospinal tract, which decussates at the lower medulla. The EEG_C3_-ESG_ipsi-anterolateral_ coherence exhibited two distinct peaks in the beta band, one in the low-beta band (13-20 Hz) and the other in the high-beta band (20-30 Hz), as shown in Figure 4a. Moreover, ESG_ispi-anterolateral_ also demonstrated significant coherence with EEG_C4_ in the low-gamma band.

**Figure 4:**
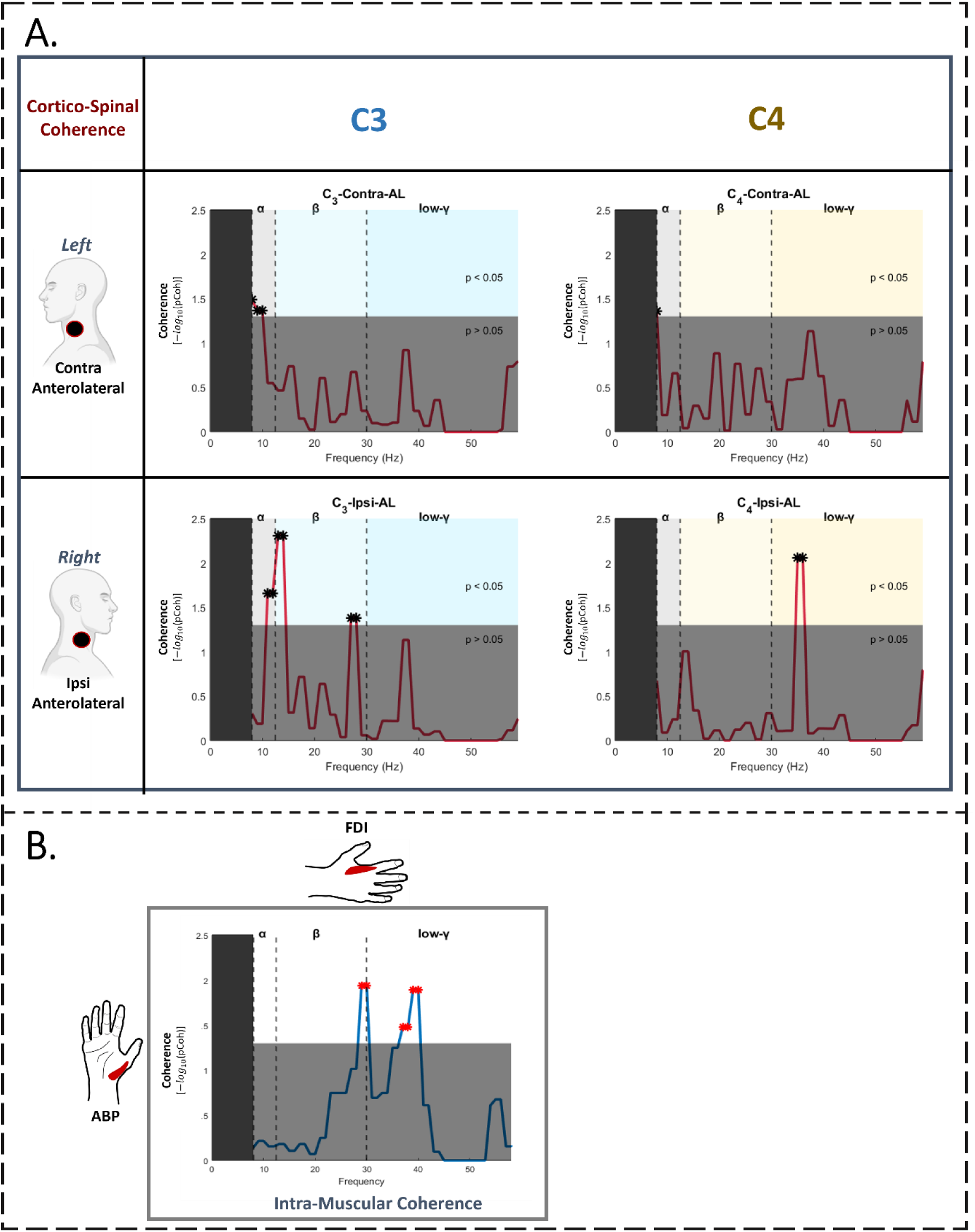
Coherence between two regions/muscles between 8Hz - 60Hz. The star indicates significant coherence at the corresponding frequency. The horizontal grey-shaded area represents non-significant coherence (p>0.05), while everything above are significant. A. Corticospinal Coherence (CSC) between two spinal regions (ipsi-anterolateral, contra-anterolateral) and two EEG signals (C3: contralateral primary motor region, C4: Ipsilateral primary motor region). Significant coherence between ipsi-anterolateral spinal region and contralateral primary cortex region (C3 EEG) was observed in the beta band, with distinct peaks at ∼13Hz-14Hz and ∼27Hz-28Hz. Ipsi-anterolateral spinal region also exhibited significant frequency-specific coherence with the ipsilateral primary motor cortex region (C4 EEG) at ∼35-36 Hz. No significant coherence was observed between the motor cortex and the contra-anterolateral spinal region. B. Inter-muscular coherence between ABP and FDI, as an indirect cortico-spinal connectivity measure. Two frequency-specific significant coherence peaks at ∼30 Hz and ∼35-40Hz were observed.

Furthermore, coherence between the Fz (frontal-midline) EEG electrode and the lower cervical regions was estimated as a control EEG-ESG connectivity condition. No significant coherence of the Fz electrode was observed with either ipsilateral or contralateral anterolateral spinal regions (S2 Figure: Control Condition: Corticospinal Coherence). The absence of significant coherence between frontal midline and anterolateral spinal regions demonstrates region-specific connectivity between motor cortex and lower cervical spinal cord, indicative of corticospinal tract connectivity.

### Indirect cortico-spinal assessment

Intermuscular coherence (IMC) between ABP and FDI was assessed as an indirect measure of corticospinal connectivity (Fisher et al., 2012). Inter-muscular coherence between ABP and FDI demonstrated significant coherence in the high-beta band (20-30 Hz) and the low-gamma band (∼40Hz).

## Discussion

To our knowledge, this is the first study to non-invasively investigate task-related spatio-spectral characteristics of spinal neural activity and its functional connectivity with the motor cortex. Our study reveals that the spinal cord exhibits frequency-band-specific spatial activation and functional connectivity patterns during isometric pincer grip hold.

### Ipsi-anterolateral beta activity, a key spinal signature

Beta activation is reported to be a dominant feature of sensorimotor pathways during voluntary motor tasks (Brown and Williams, 2005; Engel and Fries, 2010; Gilbertson et al., 2005). Increased beta-band activity during sustained contraction is widely observed in supraspinal motor regions (Baker, 2007; Chakarov et al., 2009; Kilner et al., 1999; Klostermann et al., 2007). Here, our results demonstrate a significant increase in beta-band power at the lower cervical vertebral levels (C5-C7) during the isometric hold (see Figure 2). This beta-band activity was significantly lateralised to the ipsilateral spinal region, revealing a distinct task-relevant spatio-spectral pattern of ipsi-anterolateral beta activation during sustained pincer-grip. Moreover, the relevance of the observed spatio-spectral pattern as a key signature was further highlighted by the EEG-ESG coherence, showcasing significant functional connectivity between the contralateral primary motor cortex and the ipsi-anterolateral spinal region at C5-C7 in the beta-band (see Figure 6a). These observations are consistent with the suggested characteristics of beta oscillations, including their ability to synchronise across long-distance networks (Baker, 2007; Kilavik et al., 2013; Kopell et al., 2000), and support multi-level coupling (cortex and spinal cord) via the corticospinal tract (Baker, 2007; Baker et al., 2003; Fisher et al., 2012; Jaiser, 2014).

The ipsilateral spinal region at C5-C7 vertebral levels also demonstrated significant coherence with the ipsilateral primary motor cortex in the low-gamma band (Figure 4a). This finding may reflect a complementary role of the ipsilateral motor cortex in maintaining task performance alongside the contralateral motor cortex, as proposed by previous cortical studies (Mayhew et al., 2017; Perez and Cohen, 2008; Porcaro et al., 2021; Shibuya et al., 2014). In particular, TMS-based investigations report that the ipsilateral motor cortex can influence interhemispheric inhibition and corticospinal tract output to modulate task-dependent motor activity (Davare et al., 2007; Diedrichsen et al., 2013; Perez and Cohen, 2008; Stinear et al., 2001). The observed functional connectivity between the ipsilateral spinal region and the ipsilateral motor cortex in the low-gamma band may suggest a similar task-maintenance role through direct or indirect modulation of corticospinal output.

IMC, calculated as a potential indirect measure of cortico-spinal connectivity, was comparable to previously reported IMC findings during sustained contraction (Fisher et al., 2012; Jaiser, 2014; Laine and Valero-Cuevas, 2017; McManus et al., 2019). IMC demonstrated a similar spectral profile to that of EEG-ESG-based corticospinal coherence, though it lacked spatially-relevant information. This spectral similarity supports the idea that IMC indirectly reflects corticospinal connectivity and captures task-dependent neural coupling. However, future studies are required to further investigate the neurophysiological basis of this relationship and the functional significance of such task-dependencies.

### Lateralised spinal activation, a frequency-dependent characteristic

We identified frequency-band-dependent lateralisation during the pincer-grip hold task. Although beta-band activation exhibited significant ipsilateralization (see Figure 5), no such lateralisation was observed for either low-gamma or high-gamma bands. These results could, in part, explain the conflicting results previously reported in spinal-fMRI studies during motor activity (Landelle et al., 2021). The observed contrasting degrees of lateralisation during movement using spinal-fMRI range from bilateral activity (Bouwman et al., 2008; Ng et al., 2008) to significantly lateralized activity (Maieron et al., 2007; Weber et al., 2016). Here, we observe that lateralisation is highly frequency dependent, suggesting that the variation in spectral content across different motor tasks adopted in spinal-fMRI studies could account for the observed variance in lateralisation. In particular, while beta activity is observed during sustained or isometric contractions, such oscillations are suppressed and replaced with higher frequency oscillations (gamma band) during movement (Brown, 2003; Crone et al., 1998a; Joundi et al., 2012; Miller et al., 2007; Pfurtscheller et al., 2003). Given that we observed no significant lateralisation in gamma bands, the predominant frequency content of the motor task may result in contrasting degrees of lateralisation patterns. This could potentially inform future spinal-fMRI studies to explore the effects of motor tasks and analysis methods on spinal-fMRI lateralisation and indirectly infer its effects on frequency-specific spinal activity.

**Figure 5:**
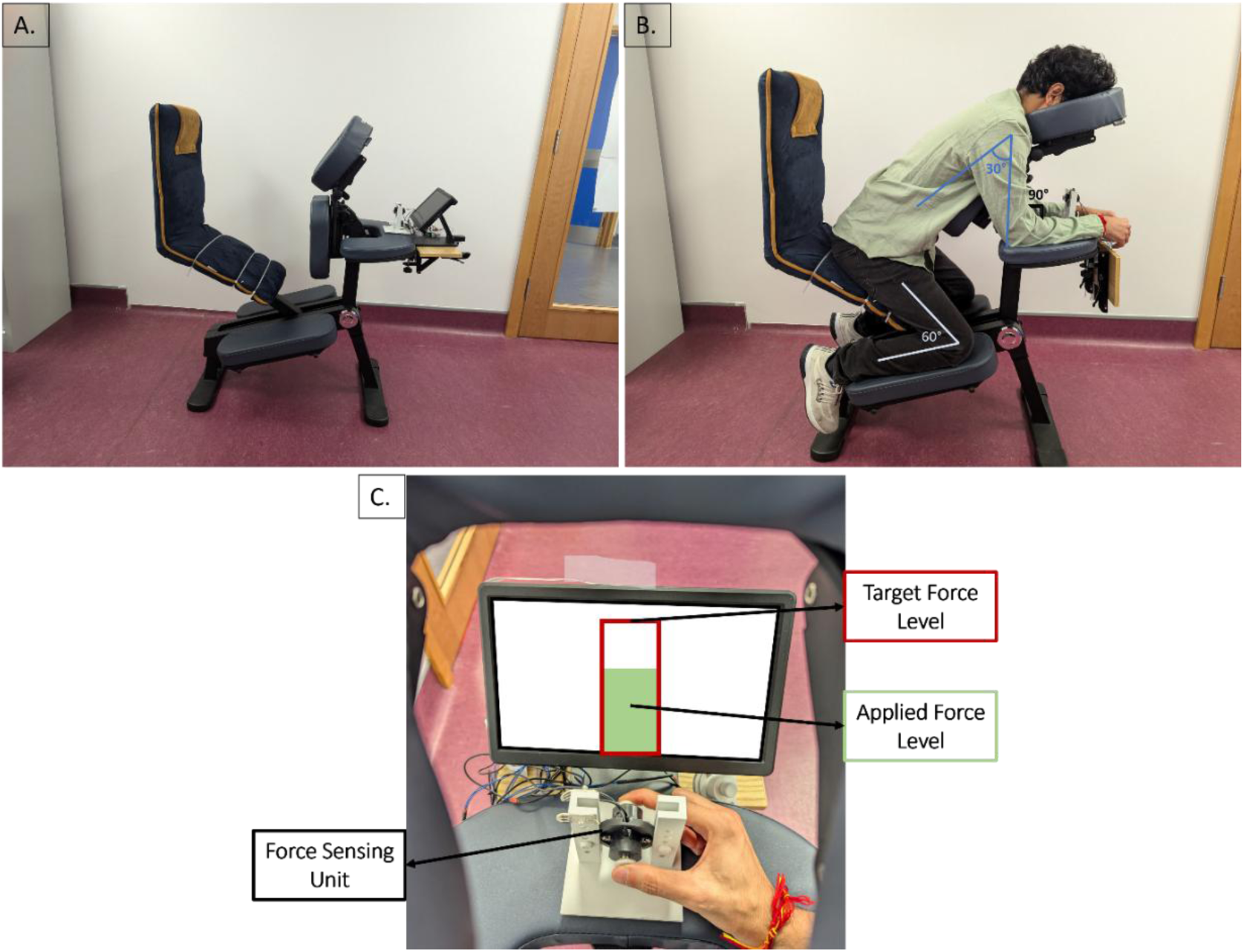
A. Experimental setup with the physio-chair and 10.1-inch screen that was used during the experiment. B. A participant sitting in an inclined position on the physio-chair. The participant’s chest was supported by the chest rest, while their head rested on the headrest, and their hands rested on the handrest, respectively. C. Visio-motor system used during the voluntary motor task and the pincer-grip hand position. The visio-motor system consists of a force sensor system and the 10.1-inch screen used for visual feedback during the task. (observed through the perspective of the participant)

Additionally, we also observed significant dorsal midline activation at C6-C7 levels in the low-gamma and high-gamma bands, which were not observed in the beta-band (see Figure 2). Supraspinal gamma-band activity (Crone et al., 1998a) has been largely associated with sensorimotor integration (Aoki et al., 1999; Ihara et al., 2003) and sensory afferent feedback (Szurhaj et al., 2006, 2005). Here, the activation of the dorsal midline might indicate dorsal horn activation, potentially corroborating gamma band activation as a signature of sensory afferent feedback and sensorimotor integration. Though this needs to be further examined in future research.

### Limitations and Future Directions

When interpreting the results of the study, some limitations should be considered. Firstly, the spatial filter used to improve the ESG signal-to-noise ratio was designed based on the evoked spinal activity recorded in response to the peripheral nerve stimulation, rather than being derived from the specific task under investigation. Secondly, the study was conducted in a relatively small and homogeneous population of ten young healthy adults, and it would be important to further validate the findings in larger and more heterogeneous populations.

Future studies will aim to address these limitations and extend the present work in several ways. The future direction of this study will aim to incorporate temporal analysis to investigate possible temporo-spatio-spectral patterns, as it is conceivable that spinal activity might demonstrate distinct temporal patterns alongside spatio-spectral signatures during the pincer-grip task. Furthermore, the connectivity analysis will be extended to investigate spino-muscular connectivity (ESG-EMG coherence). Spino-muscular connectivity, in conjunction with cortico-spinal connectivity investigated here, can reveal connectivity patterns at different levels along the corticomuscular pathway. Finally, investigation of EEG-ESG-based cortico-spinal connectivity in individuals with neurological conditions could reveal patterns of corticospinal dysfunction.

## Conclusion

In conclusion, our study revealed that the spinal cord exhibits distinct frequency-specific spatial activation and cortical connectivity patterns during sustained pincer-grip hold. Notably, the ipsi-anterolateral spinal region demonstrated high task-relevance during sustained hold. Ipsi-anterolateral spinal region exhibited significant activation and connectivity with the primary motor cortex in the beta-band, potentially indicating neural activity from the ventral grey matter (anterior horn), which features motor-neuronal links with both supraspinal systems and skeletal muscles (Haines et al., 2018). Similar to its cortical counterpart, the beta band was observed to be a key spectral signature of spinal cord activity at lower cervical levels, further underscoring its relevance during sustained control.

## Materials and Methods

### Participant and Ethical approval

Ten healthy participants (n=10, age: 28 ± 4, 4 Males) were recruited for the study. All participants were informed about the experiment paradigm, and signed consent was obtained before the session. The ethics was approved by ‘Tallaght Hospital/St. James’s Hospital Joint Research Ethics Committee, Project Reference: 0493, and the study was performed in accordance with the declaration of Helsinki.

### Participant Exclusion Criteria

The exclusion criteria are as follows:

1. No history of neuromuscular, neurological, and or psychiatric condition<u>s</u>.
2. No diagnosed chronic cervical and upper back condition (affecting spinal nerves, and or muscles).
3. No history of spondylosis or spondylitis or any other back pain / spinal degeneration.

### Experimental Paradigm

This observational study recorded multimodal signals during the experimental paradigm of the voluntary motor task followed by peripheral nerve stimulation. During the voluntary task, participants were instructed to keep their eyes open. However, during the peripheral nerve stimulation, they were instructed to keep their eyes closed. The stimulation paradigm was included with the objective of designing a spatial filter to improve the signal-to-noise ratio of ESG signals recorded during the motor task. The results from stimulation paradigm are reported in the Appendix/Supplementary Material (S1 Figure).

#### Voluntary Motor Task

Each participant performed an isometric pincer-grip contraction during the voluntary task paradigm. The task was performed using a force sensor block held between the thumb and the index finger of their right hand. During this experimental paradigm, the participant performed 30 sessions of the visually guided voluntary task. Each session lasted approximately 15s and consisted of three guided phases: rest phase (5s), isometric contraction phase (5s), and relaxation phase (5s) Figure 1. During the isometric contraction phase, the participant maintained a pincer grip force at 10% of their maximal voluntary contraction (MVC). This was visually guided by displaying fixed target force and real-time applied-force feedback on the screen (Figure 5c). The MVC was determined at the start of the paradigm as the average peak force during three short maximal contraction trials (3s each) with a 30s inter-trial rest period.

#### Peripheral Nerve Stimulation

The peripheral nerve stimulation paradigm consisted of 35 sessions of median nerve stimulation performed on the wrist at an intensity of 1.5 times the motor threshold and 200 µs pulse width. Motor threshold was characterised as the minimum intensity at which thumb movement was observed. Each session lasted for 30 s; 20 s of the stimulation phase followed by 10 s of the rest phase. During the stimulation phase, the median nerve was stimulated with a train of pulses at an average pulse frequency of 2 Hz, resulting in 40 stimulation responses per session. A total of 1400 responses (35 sessions x 40 trials per block) to median nerve stimulation were recorded.

### Experiment Setup

The participant sat comfortably in a physio-chair for the duration of the experiment, as shown in Figure 5a. The participant’s head comfortably rested on the headrest of the physio-chair and the hand rested on the handrest while making a 90-degree angle at the elbow (Figure 5c).

Peripheral Nerve Stimulation: A MDD CE medically certified constant current stimulator, DS7A, was used for median nerve stimulation at the wrist.

### Data Acquisition

A total of 210 channels of data were recorded simultaneously using the high-speed Biosemi-ActiveTwo System (Biosemi B.V., Amsterdam, The Netherlands) at a sampling frequency of 8192 Hz. This system was employed to collect high-density EEG, high-resolution ESG, bi-polar EMG, and ECG signals, as shown in Figure 6. Visual inspection was conducted on all EEG and ESG channels; electrode offset was maintained between ± 20mV and ± 25mV, respectively.

**Figure 6:**
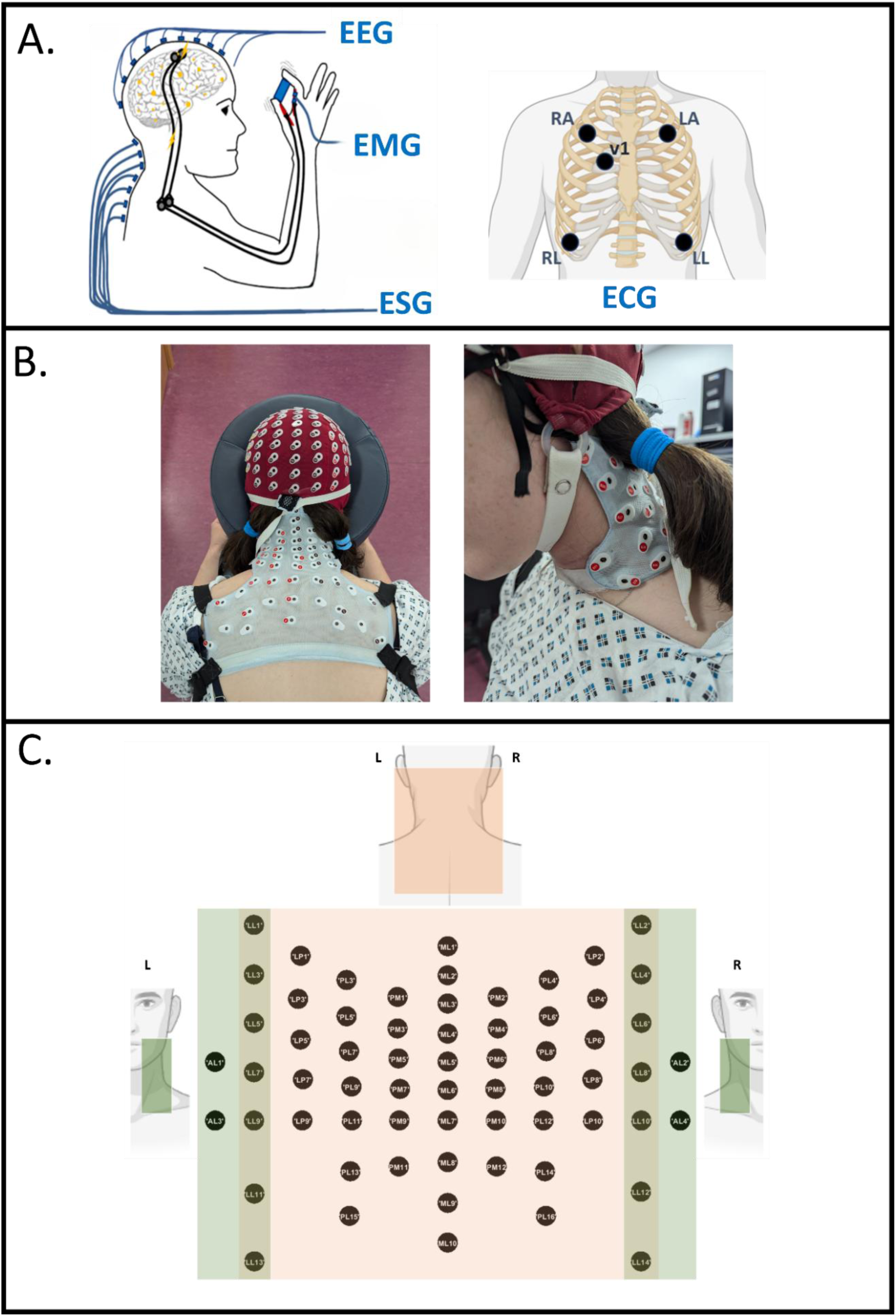
Schematic representation of electrophysiological data collected during the experiment. B. The HD-EEG and HD-ESG electrodes positioned over the participant’s head, neck, and upper back, highlighting the electrode locations used for recording the HD-EEG and HD-ESG signals during the experiment paradigms. C. 64-channel derived from the SC10X/U system utilised for recording HD-ESG.

#### Electroencephalography (EEG)

Signals were recorded using a 128-channel high-density active electrode system. The EEG electrodes were derived from the 10-5 electrode system (Biosemi B.V., Amsterdam, The Netherlands).

#### Electrospinography (ESG)

Signals were recorded using a 64-channel high-density active electrode system. The ESG electrode placement followed the SC10X/U electrode system, as shown in Figure 6c (Mehra et al., 2025).

#### Electromyography (EMG)

Bipolar-surface EMG was recorded from two muscles: first dorsal interosseous (FDI) and abductor pollicis brevis (APB). A unipolar reference electrode was located on the bony prominence at the wrist. Signals were recorded using flat active sintered Ag-AgCl electrodes. Each electrode comprised a circular recording area (d=3 mm) within a 17x10 mm casing.

Electrocardiography: Cardiac activity was recorded in accordance with 5-electrode ECG system using flat active sintered Ag-AgCl electrodes. The 7-lead ECG signals were extracted using the 5-electrode ECG system.

1. *Lead I = LA − RA*
2. *Lead II = LA − RA*
3. *Lead III = LA − RA*
4. 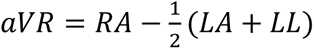
5. 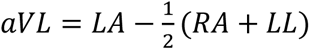
6. 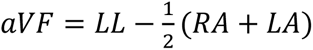
7. 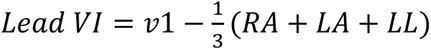

### Signal Processing

#### ESG Processing

##### I. Pre-Processing

The stimulation artifact recorded during the peripheral stimulation paradigm lasted 1-2 ms after the stimulation trigger and was removed by interpolating (piecewise cubic interpolation) the signal between 0.5 ms before the stimulation trigger and 2.5 ms after the stimulation.

The ESG signals recorded during both paradigms were high-pass filtered above 1Hz (dual-pass Butterworth filter). The ESG channels were visually inspected, and bad channels demonstrating excessive noise, persistent artefacts or intermittent signal dropout were rejected. The ECG artifacts observed in the recorded ESG signals were removed using canonical correlation analysis (CCA) (de Cheveigné et al., 2018; Hotelling, 1936). To obtain the ECG artifact relevant components, good ESG channels and the 7 ECG-leads were bandpass filtered between 5 Hz and 45 Hz (dual-pass Butterworth filter) and subjected to CCA. Canonical components with a correlation greater than 0.8 between ESG and ECG were marked. The high-passed ESG (>1Hz) was transformed to CCA space, and the marked canonical components were removed. The remaining components were back-transformed to the original ESG space, resulting in reconstructed ESG signals with little to no ECG artifact. Before further processing of the ESG signals, bad channels were interpolated based on the weighted average method using neighbouring channels (neighbour distance <= 50mm). The signals obtained after artifact rejection and bad channel interpolation were re-referenced using ‘common average referencing’. A zero-phase comb-notch filter was then used to remove 50Hz harmonic line noise.

The pre-processed ESG signals during the voluntary motor task were segmented into 30 trials of 15 seconds: 5 seconds before the start of the contraction phase and 5 seconds after the end of the contraction phase (-5s to 10s). The pre-processed ESG signals during the peripheral stimulation paradigm were segmented into 1400 trials of 0.5 seconds: 0.1 seconds before stimulation and 0.4 seconds after stimulation.

The resulting trials for each paradigm were visually inspected for bad trials. The identified bad trials for each paradigm were rejected. The remaining trials (voluntary task: 26.6 ± 1.7, ∼88%; stimulation task: 1287 ± 91 trials, ∼92 %) were bandpass filtered between 2Hz - 250Hz and 10Hz - 1400Hz (Mehra et al., 2025) for voluntary task ESG and peripheral stimulation ESG signals respectively.

##### II. Spatial Filtering

The pre-processed trials during the peripheral stimulation paradigm were subjected to task-related component analysis (TRCA) (Nakanishi et al., 2018; Tanaka et al., 2013). TRCA was proposed by Tanaka et. al. to extract the task-related components that maximise the covariances between trials within a specified time window. TRCA computes spatial filters by solving a generalised eigenvalue problem, where the goal is to find spatial projections that maximise the ratio of inter-trial covariance to overall signal variance. In this formulation, the eigenvalues quantify the degree of trial-to-trial consistency, and the corresponding eigenvectors define the spatial filters that extract reproducible task-related components. This results in increased consistency and reproducibility across trials. A 15 ms time window (7-22 ms post-stimulation) was selected for designing the TRCA-based spatial filter to maximise the inter-trial consistency of the expected early spinal evoked responses.

The significant task-related components were selected by comparing the resultant eigenvalues against the eigenvalues obtained via the null-hypothesis. The null hypothesis involved calculating eigenvalues using randomised 15 ms time-blocks sampled uniformly across the entire trial duration (Tanaka et al., 2013). This was repeated three times to obtain a distribution of eigenvalues under the null hypothesis. Task-related components with eigenvalues outside the 95% confidence interval of the null distribution were chosen as the statistically significant components.

The significant components were back-transformed to the original ESG space, resulting in a TRCA-spatial filtered signal. TRCA-spatial filtering resulted in an improved signal-to-noise ratio where the task-relevant spinal signals were retained, and the noise was removed. The trials during the voluntary motor task paradigms were also spatially filtered using the TRCA spatial filter obtained using the evoked spinal signals during the stimulation paradigm.

The signals obtained after TRCA-spatial filtering for both paradigms were re-referenced using the surface Laplacian method (Hjorth, 1975), further improving the spatial resolution of the signals prior to analysis.

#### EEG Processing

The 128-channel EEG signals recorded during the voluntary motor task were bandpass filtered between 1 Hz – 100 Hz (dual-pass Butterworth filter). The bad channels were visually identified and removed. ICA was applied to the remaining EEG signals. The independent component (IC) visually related to the blink artifact was removed and the remaining ICs were back-transformed to the original EEG space, resulting in blink artifact rejected EEG signals. The bad channels were interpolated using the weighted average of neighbouring channels. The signals obtained after artifact rejection and bad channel interpolation were re-referenced using ‘common average referencing’. A zero-phase comb-notch filter was then used to remove 50Hz harmonic line noise. 30 epochs (trials) of 15 s were selected from the resulting EEG signals, starting 5 s before the start of the contraction phase till 5 s after the end of the contraction phase.

The epoched signals were resampled at 1000Hz (anti-aliasing). ‘C3’, ‘C4’, and ‘Fz’ channels (10-5 system) were selected for further processing. The selected channels were band-pass filtered between 2-125 Hz (dual-pass Butterworth) and re-referenced using surface Laplacian (Hjorth, 1975). The signals from the resulting channels were then divided into 1 s epochs with 50% overlap during the sustained phase, between 0.75 and 4.75 seconds after the visual cue. This provided four 1 s epochs for each good trial. All 1s epochs were subjected to automated artifact rejection routines (Fieldtrip Toolbox) (Oostenveld et al., 2011) to discard contaminated data.

#### EMG Processing

The bipolar EMG signals recorded from the APB and FDI were baseline-corrected and bandpass-filtered between 10-500Hz (dual-pass Butterworth filter). A zero-phase comb-notch filter was then used to remove 50Hz harmonic line noise. The resulting signals were divided into 1 s epochs, with 50% during the sustained phase, between 0.75 s and 4.75 s after the start of the contraction visual cue. The resulting epochs were resampled at 1000Hz.

### Data Analysis

#### Task-Related Analysis

For task-related analysis, processed ESG signals during the voluntary task were resampled at 1000 Hz. The resulting signals during the sustained contraction phase (0.75s - 4.75s post-contraction cue) were segmented into Hanning windowed 1 s epochs with 50% overlap (Bayram et al., 2023; Polat and Güneş, 2007).

##### Spectral Analysis

The spectral power was calculated at integer frequency values between 5 Hz and 95 Hz using Welch’s method, for each epoch. The average spectral power across all epochs for each participant was computed in three frequency bands: beta band (13 – 30 Hz), low gamma band (31 – 60 Hz), and high gamma band (61 – 90 Hz). For each band, the average band-specific baseline power was computed for the 2 s time window, between 4.5 s and 2.5 s before the contraction cue (baseline time window: -4.5 s to -2.5 s).

Band-specific ESG activation during the sustained task was examined by identifying the channels demonstrating statistically different band-specific power during the task compared to baseline. For each band, this resulted in a set of channels exhibiting task-relevant ESG activation, and the baseline-corrected spectral power calculated for these channels was defined as task-relevant power (pow_task_). Topographical maps of the resulting band-specific task-relevant power were plotted over the SC10X/U electrodes. To assess activity in the ipsi-anterolateral ESG region, channels LL8 and AL4 were selected, similarly, channels LL7 and AL3 were selected to assess activity in the contra-anterolateral ESG region. For each frequency band, regional activity was quantified as the percentage change in spectral power during the task relative to rest, for the selected electrodes. Band-specific lateralisation was evaluated by calculating significant differences in the regional activity between the two regions.

##### Statistics

For determining band-specific task-relevant power, the Wilcoxon signed-rank non-parametric test was applied at each channel to identify statistically significant spectral power (task compared to baseline) for each band. Adaptive false discovery rate (FDR) at q = 0.05 was used to correct for multiple comparisons (Benjamini et al., 2006). For determining band-specific lateralisation, significant differences between the regions for each band were determined using the Wilcoxon signed-rank non-parametric test.

##### Coherence Analysis

Cortico-spinal coherence between EEG and ESG signals during sustained contraction was calculated using the method described by Bista et. al. (Bista et al., 2023; Coffey et al., 2021). For this analysis, two EEG channels (C3 and C4) and two ESG regions, ipsi-anterolateral (LL8 and AL4) and contra-anterolateral (LL7 and AL3), were selected. The C3 and C4 channels were chosen to examine cortical activity over the contralateral and ipsilateral primary motor regions, respectively.

Coherence analysis was applied to 1-second EEG and ESG epochs. One participant was excluded from analysis due to poor EEG signal quality, with more than 10% of total EEG channels identified as bad. For each remaining participant, coherence was estimated for all combinations of EEG channels and ESG regions.

The cross-spectrum for a given EEG–ESG region pair was estimated by first computing the cross-spectra between the EEG channel and each ESG channel within the region, then averaging these cross-spectra. The auto-spectrum for each ESG region was computed by averaging the auto-spectra of all ESG channels within that region (e.g., LL8 and AL4 for the ipsi-anterolateral region). Both auto-spectra and cross-spectra were smoothed using a spatial median at a 2-Hz frequency resolution. Finally, the smoothed cross-spectra were normalised with respect to the corresponding auto-spectra to estimate 2-Hz banded cortico-spinal coherence (Bista et al., 2023; Coffey et al., 2021).

Likewise, inter-muscular coherence (IMC) was estimated between bi-polar FDI and ABP EMG, as an indirect marker of cortico-spinal connectivity (Fisher et al., 2012). Additionally, coherence between the Fz EEG channel and the two ESG regions was estimated as a control for the cortico-spinal coherence analysis.

##### Statistics

The non-parametric one-sample spatial signed-rank test for spectral coherence (Nasseroleslami et al., 2019; Oja, 2010; Oja and Randles, 2004) was applied to compare each coherence value against zero, resulting in individual p-values for each band per channel-combination per participant. The respective p-values represented the strength of the normalised coherence. Stouffer’s method (Stouffer et al., 1949; Westfall et al., 2014) was used to combine individual p-values across all participants to derive group-level average p-values. Adaptive false discovery rate (FDR) at q = 0.05 was used to correct for multiple comparisons (Benjamini et al., 2006). The resulting group-level strength (p-value) for each coherence value per combination was visualised by taking the negative log of the p-values.

## Appendix/Supplementary Material

### S1 Text: Evoked Response Analysis

The processed ESG signals during the peripheral nerve stimulation paradigm were averaged across all epochs, and the latency characteristics of the resulting evoked spinal response potentials were evaluated.

The evoked ESG responses were further averaged over all the participants, resulting in grand-mean evoked responses for the participant population. The grand-mean evoked activity between 12 ms and 15 ms was mapped across the SC10X/U electrode space to analyse the spatial distribution of the evoked spinal activity. The results from this analysis are reported as supplementary material (S1 Figure).

#### Statistics

Statistically significant evoked response was determined at each sample point of the evoked response waveform for each channel. The significance was determined by comparing the difference of the evoked amplitude from zero across all participants, using the Wilcoxon signed-rank non-parametric test (Iyer et al., 2017; Nasseroleslami et al., 2014). To account for multiple comparisons, PCA was applied between -0.001s and – 0.03s time windows across 64 channels and the Wilcoxon signed-rank test was applied to the reconstructed ESG signal that retained dominant principal components explaining 85% of the total variance, plus an additional component. Bonferroni correction was subsequently applied at α = 0.05.

**S1 Figure:**
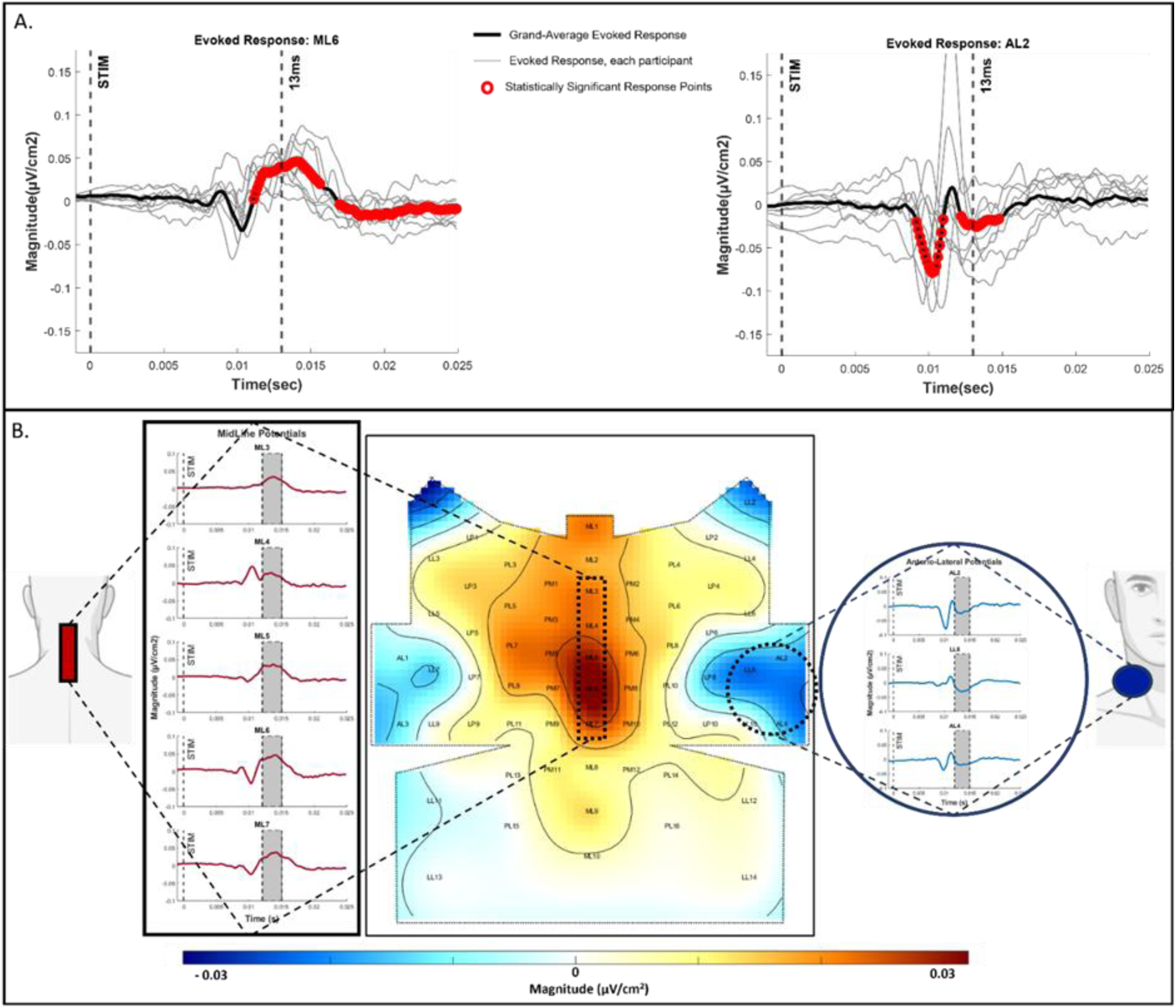
A. Grand-mean of the evoked potentials across all participants at ML6 and AL2 electrode positions. Significant Evoked potentials: P9 (mean ± sd), and N13/P13 (13.2 ± 1.1ms) were observed in response to the median nerve stimulation at these electrode positions. Grey lines represent individual participant evoked potentials, and thick black line correspond to the grand mean evoked potential across all participants. The statistically significant evoked response sample points are illustrated as hollow red circles. B. Topographic Map of Evoked Spinal Potentials: Centre Figure: Topographic Map of the grand-mean spinal evoked response over the SC10-X/U sensor space, time-window (12ms-15ms). Left figure: corresponds to the spinal activity recorded from channels ML3-ML7 (midline, posterior) for the time frame between 0.001s before stimulation and 0.025s after stimulation. Right figure corresponds to the spinal activity recorded from lateral-anterior channels for the time frame between 0.001s before stimulation and 0.025s after stimulation. This figure demonstrates the epicentre of activity at dorsal channels ML5-ML6 and a polarity reversal of this activity at ipsi-anterolateral channels (LL8, AL2, AL4).

**S2 Figure:**
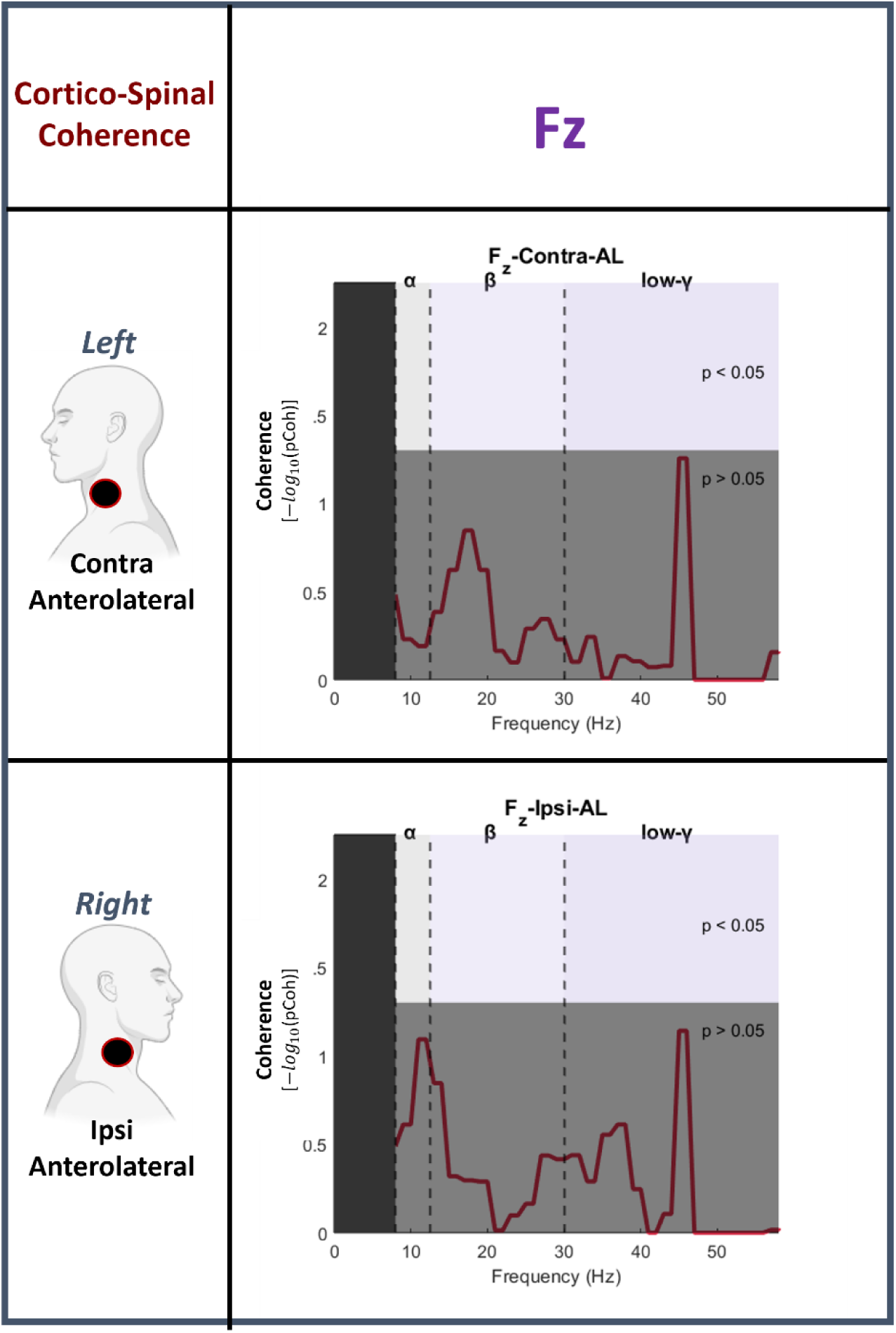
Corticospinal Coherence (CSC) between the Fz EEG electrode and two spinal regions (ipsi-anterolateral, and contra-anterolateral). Fz demonstrated no significant coherence with either contra or ipsi-anterolateral spinal region, in the frequency range of 8Hz – 60Hz. The grey-shaded area represents non-significant coherence values (p>0.05).

